# Encapsulation in a bacterial microcompartment shell improves thermal stability of a glycolytic enzyme

**DOI:** 10.64898/2026.01.28.702358

**Authors:** Nicholas M. Tefft, Neetu S. Yadav, Megan C. Gruenberg Cross, Charles D. Swiggett, Kristin N. Parent, Josh V. Vermaas, Michaela A. TerAvest

## Abstract

Selective encapsulation of target enzymes is an increasingly well studied field with a host of potential applications for biotechnology. Natively, many bacteria utilize bacterial microcompartments (BMCs) for enzyme encapsulation to enhance catalysis. BMCs are protein shells that enable selective localization of targeted metabolic enzymes and may improve catalytic rates by co-localizing pathway enzymes and/or serve to sequester toxic or volatile intermediates. The microcompartment shell of *Haliangium ochraceum* (HO) is a notable BMC chassis because of its modularity and versatility; it is easily expressed and assembled outside its native host and can accept a wide array of cargo. Recently, it was demonstrated that assembly of HO BMC shells can be easily achieved *in vitro*. Following up on our previous work on *in vivo* assembly of HO-BMCs with triose phosphate isomerase (TPI) as model enzyme cargo, here we have demonstrated the advantages of *in vitro* assembly (IVA) for targeted enzyme encapsulation. We achieved variable loading of BMC shells with targeted amounts of TPI and demonstrated enhanced thermal stability of encapsulated TPI versus free TPI up to 62°C.

## Introduction

Bacterial microcompartments (BMCs) occur in many bacteria and encapsulate a range of different metabolic pathways, including carbon dioxide fixation and propanediol catabolism.^1–5^ BMCs are polyhedral protein shells that localize target proteins and create internal environments that enhance catalysis and/or sequester toxic intermediates.^6–9^ Native BMC functions have been studied in several systems, with carbon-fixing carboxysomes being particularly well studied.^10–13^ BMCs may also be useful for pathway engineering and the BMC shell from *Haliangium ochraceum* (HO) has been repurposed as a versatile platform for engineered encapsulation. Target proteins can be easily and rapidly confined in HO BMC shells to take advantage of the benefits of encapsulation for engineered metabolic pathways.^14,15^ The HO BMC shells used in this platform are assembled using the following protein oligomers: a hexamer (BMC-H), three different trimers (BMC-T1, -T2, and -T3) and a pentamer (BMC-P).^16–18^ Four distinct icosahedral BMCs can be formed from different combinations of these subunits. Minimal shells (HT1) use only the monolayered trimer, T1, while full shells (HT1T2T3) also incorporate the stacked BMC T2 and T3 trimers. Minimal and full shells may both be made in capped (including the BMC P pentamer) or uncapped, ‘wiffle ball’ (excluding BMC P) versions.^15^ Wiffle ball shells have large gaps where the pentamer is absent from the vertices. The engineered SpyCatcher-SpyTag system for protein-protein covalent linkage has been used in several instances to localize non-native cargo to HO BMC shells.^14,19–21^ This has typically been achieved by fusing the SpyTag peptide to T1 and the SpyCatcher domain to the desired cargo(s).

We previously utilized the SpyCatcher-SpyTag system^14,22^ to localize triose phosphate isomerase (TPI), into all four of engineered HO shell types using *in vivo* assembly.^23^ In this method, SpyTag was appended to the T1 sequence, SpyCatcher was appended to the cargo sequence, and all shell proteins and the cargo were co-expressed in *E. coli* and purified as assembled, cargo loaded BMC shells. Little to no difference in TPI activity was observed between shells with and without pentamers, indicating that the small pores in the shell tiles enable substrate and product transport. This was supported by modeling that showed HO BMC shells (whether capped or uncapped) generally are not a significant barrier to diffusion of small molecules.^24,25^ Full shells with all trimer types had significantly lower TPI activity due to fewer available SpyTags to localize the cargo (because only T1 and not T2 nor T3 contained the SpyTag fusion). While using varying tile combinations enabled some control of cargo loading, the *in vivo* assembly method limited our ability to finely control cargo levels within the shells. Further, proteomic analysis indicated contamination of native proteins in HO BMC shells, which may have resulted in reduced cargo loading and caused background TPI activity in ‘empty’ shells. These results indicated the disadvantages of assembling and loading the shells *in vivo*, where there are competing cargo molecules and a low level of control over cargo concentration.

Recent work demonstrated that *in vitro* assembly (IVA) is a promising route for further development of the HO BMC platform.^26^ Multiple strategies for IVA of the HO BMC have been demonstrated, but there are several key advantages of the recently published chaotrope-based method.^27^ In this procedure, BMC shell components and cargo are combined in a stepwise manner, with BMC-H added last. BMC-H solutions for assembly contain high urea concentrations (8 M) to block the previously documented self-assembly of BMC-H into sheets and tubes ^28^. Dilution of the BMC-H solution into the IVA mixture containing other cargo and shell components triggers rapid shell assembly by a sudden reduction in urea concentration. Stepwise addition also allows cargo molecules to conjugate to the trimer before the shell is fully assembled. Notably, the speed of assembly using this method is fast, with full formation of shells in under 2 minutes.^26^ IVA is also advantageous because large amounts of each subunit and cargo can be easily purified separately, and assembly can be scaled based on the application. This also enables iterative assemblies to be carried out rapidly with small changes to shell component ratios. Similarly, co-conjugation of multiple cargos will likely be simple using this IVA method, although this needs to be tested. These advancements enabled significant improvements to assembly of BMCs containing TPI and iterative testing of varying conditions.

One advantage of BMCs as a protein engineering platform is that they are very stable and retain their structure at increased temperatures, low and high pH, and over long time scales.^29,30^ It may be reasonable to expect that proteins encapsulated within the shells may be similarly protected. Proteins rely on proper folding to function and while increasing temperature generally increases the rate of reaction for enzymes, there is an impact on three-dimensional structures at higher temperatures that can cause denaturation.^31,32^ Many enzymes used for biotechnology are derived from mesophilic bacteria,^33^ including the TPI used in this study, which is from *E. coli* (Table 1). However, industrial processes can benefit from operating at thermophilic conditions to enhance reaction rates and reduce contamination.^33–35^ Improving thermal stability of proteins through modification or identification of thermophilically stable homologs is highly sought after.^34,36–38^ However, not all potential catalytic targets are easily modifiable or can be found in thermophiles. Here, we hypothesized that encapsulation in HO BMC shells could enhance thermal stability of enzymes. Our previous work using TPI as a model enzyme is well positioned to investigate thermal stability of encapsulated enzymes.^23^ In this work, we investigated the advantages of *in vitro* assembly (IVA) and used these methods to test if BMCs confer thermostability on encapsulated cargo.

**Table 1.**
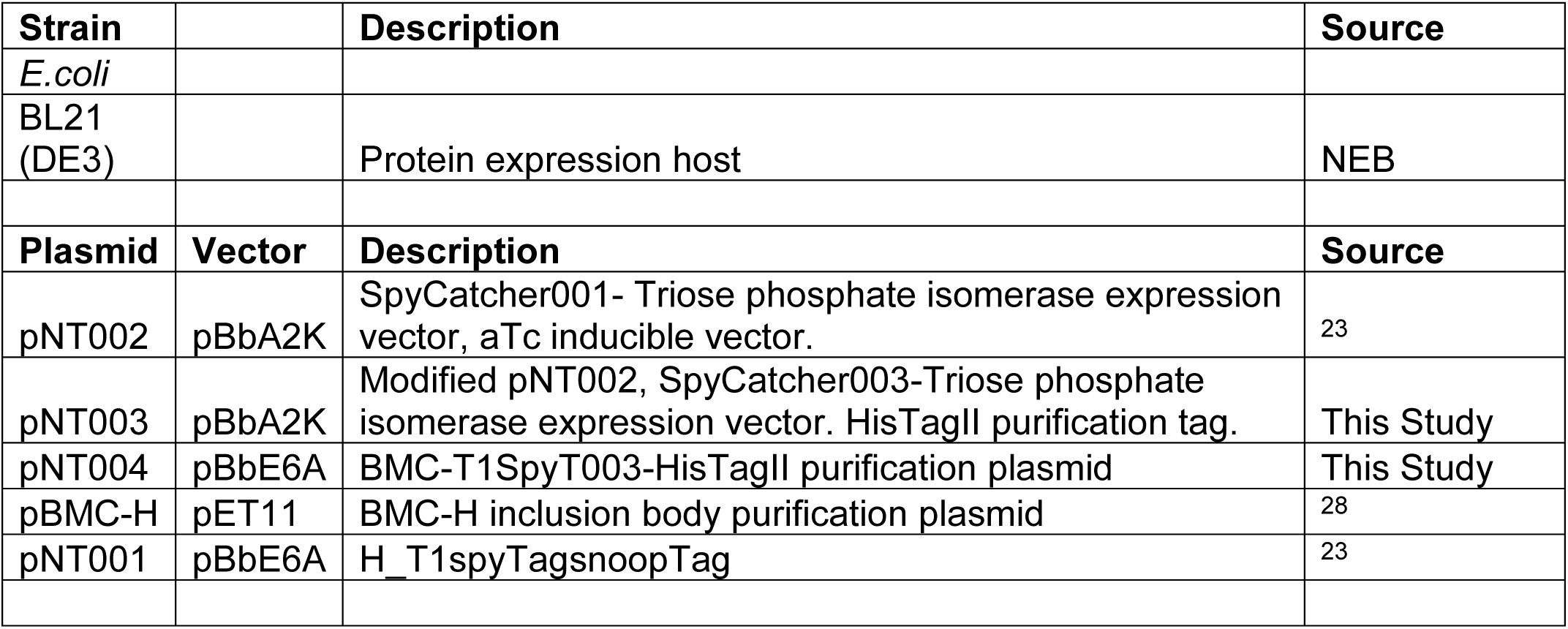
Strains and plasmids used in this study.

| Strain |  | Description | Source |
| --- | --- | --- | --- |
| *E. coli* |  |  |  |
| BL21 (DE3) |  | Protein expression host | NEB |
| Plasmid | Vector | Description | Source |
| pNT002 | pBbA2K | SpyCatcher001- Triose phosphate isomerase expression vector, aTc inducible vector. | 23 |
| pNT003 | pBbA2K | Modified pNT002, SpyCatcher003-Triose phosphate isomerase expression vector. HisTagII purification tag. | This Study |
| pNT004 | pBbE6A | BMC-T1SpyT003-HisTagII purification plasmid | This Study |
| pBMC-H | pET11 | BMC-H inclusion body purification plasmid | 28 |
| pNT001 | pBbE6A | H_T1spyTagsnoopTag | 23 |

## Results

### Design and purification of TPI-loaded HO shells via *in vitro* assembly

In this study, we used a chaotrope-based approach for *in vitro* assembly of HO BMC shells, which was recently developed by Range et al.^26^ In this method, buffer, trimer(s), the target cargo are combined first, then the urea-solubilized hexamer is added. In our previous work, we used SpyCatcher001 and SpyTag001 sequences for conjugation. We found that conjugation of TPI to T1 was slow and caused sub-optimal shell loading with the original fusion sequences. We solved the slow conjugation by upgrading the fusions to SpyCatcher and SpyTag version 003 sequences, which show up to 400-fold faster conjugation rates.^39^ With the new sequences, assembly of shells with activity comparable to that in our previous work was attainable in as little as fifteen minutes.

We first compared HT1 TPI shells produced by *in vitro* assembly with HT1 TPI shells produced using *in vivo* assembly.^23^ The *in vivo* produced shells were estimated to have 1 TPI conjugated to each trimer tile and therefore, were compared to IVA-produced HT1 TPI shells that were assembled with an expected cargo ratio of 1 TPI dimer per T1 tile. We used a commercial TPI assay kit to compare activity of various shell samples. The activity of IVA-produced HT1 TPI was significantly higher than the *in vivo* produced shells per mg total protein (p = 0.03, student’s *t*-test) suggesting IVA shells achieved higher TPI loading than previously observed (**Figure 1**), which may be attributable to fewer off target proteins being encapsulated, allowing more TPI to be loaded. The TPI activity of *in vivo* generated HT1 shells (‘no cargo’) was significantly above background, as previously observed, likely due to adventitious capture of native TPI. However, IVA HT1 shells showed nearly zero activity, indicating that switching to IVA reduced the capture of unwanted proteins inside the BMC shells.

**Figure 1.**
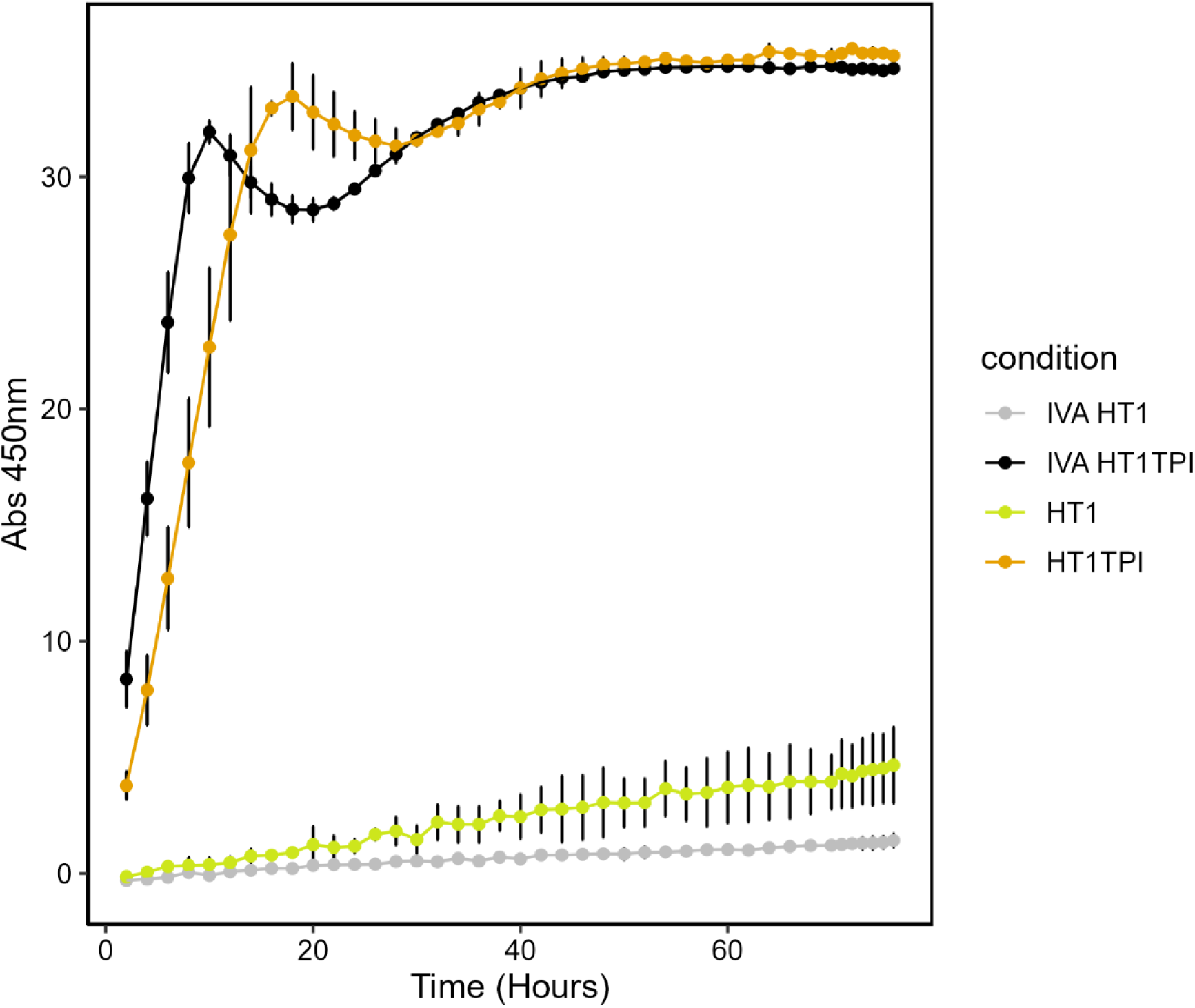
Tpi Activity comparison of *in vitro* and *in vivo* HO shells. Increase in absorbance at 450 nm (A_450_) over time in a Tpi activity assay with 238 ng protein loaded per well for each sample. Grey is IVA HT1, yellow is *in vivo* HT1, black is IVA HT1 TPI, orange is *in vivo* HT1 TPI. Each point represents the average of two replicates with standard deviation shown in error bars.

### Targeted loading of variable concentrations of triose phosphate isomerase cargo

With dependable *in vitro* cargo conjugation and shell assembly, we achieved variable loading of shells by increasing or decreasing available TPI in the assembly reaction. We assembled with the following T1:TPI ratios: 6:1, 3:1, 3:2. 1:1, 1:2, 1:3. These ratios are expressed as T1 trimer:TPI dimer, i.e., 1:1 indicates one TPI dimer per trimer tile. We observed clear differences in TPI-T1 conjugate and free T1 abundance on SDS-PAGE across TPI ratios (**Figure 2A**). Specifically, the band for free T1 became less intense, and the band for The T1-TPI conjugate became more intense as more TPI-SpyCatcher was added to the reactions. To evaluate differences in cargo loading, TPI activity was measured, and results were consistent with the expected loading (**Figure 2B**). Shell assemblies with T1:TPI ratios of 6:1, 3:1, and 3:2 showed lower activity than 1:1, which was predicted to have 1 TPI dimer per trimer tile. Similarly, TPI loading ratios of 2 and 3 per trimer showed higher activity than TPI 1, although the increase in activity at higher loading begins to saturate.

**Figure 2.**
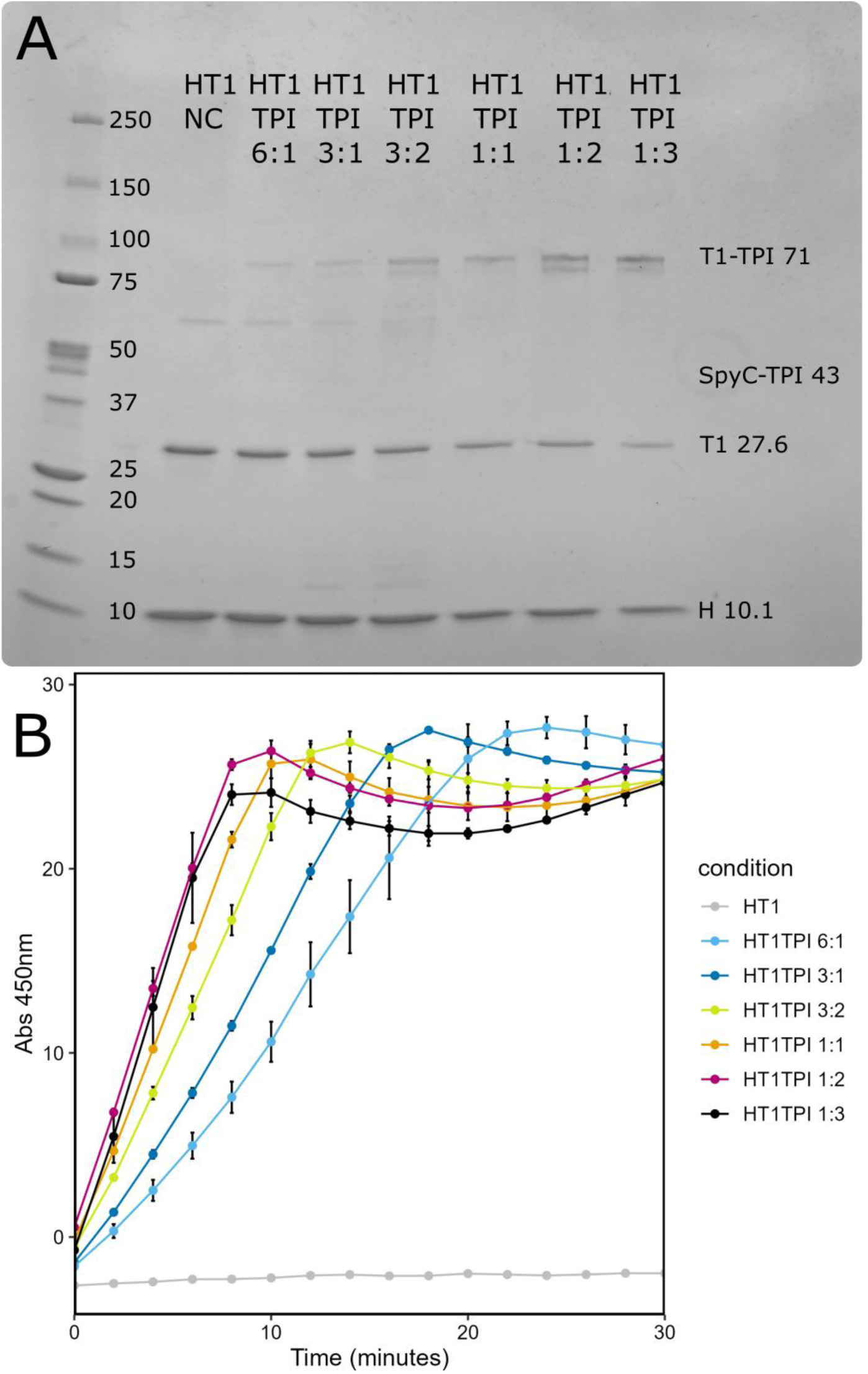
**A. SDS-Page of variably loaded TPI HO shells** Coomassie strained SDS-PAGE, 480 ng of protein loaded per well. **B. Tpi Activity of variably loaded HO shells.** Increase in absorbance at 450 nm (A_450_) over time in a Tpi activity assay with 238 ng protein loaded per well for each sample. Grey is HT1, Light blue is HT1 TPI 6:1, dark blue is HT1 TPI 3:1, yellow is HT1 TPI 3:2, orange is HT1 TPI 1:1, magenta is HT1 TPI 1:2, black is HT1 TPI 1:3. Each point represents the average of two replicates with standard deviation shown in error bar.

Shell assembly was first evaluated using dynamic light scattering (DLS) (**Table S1**) and the majority of shells measured between 44 and 46 nm, which falls within the expected size of HO-BMCs. This result indicated that shells assembled as expected and was further supported by the UV-vis traces taken from fast protein liquid chromatography (FPLC) during separation using the size exclusion chromatography (SEC) column. SEC traces showed peak areas for the void volume that were consistent with expected yield for each assembly ratio (**Figure S1).**

SEC traces also showed that the formation of T1-TPI conjugates that were not included in shell assembly also increased as the TPI ratio increased, this is likely due to a combination of assembly time and steric hindrance preventing additional encapsulation. These results suggested shells assembled normally, however DLS showed a trend of increasing diameters in the shells with more TPI loading. HT1 TPI 1:1 measured at 47 nm ± 0.11, 1:2 at 49 ± 0.21 nm, and 1:3 at 58 ± 1.00 nm. The large jump in the diameter of the HT1 TPI 1:3 sample suggested aggregation and cryo-electron microscopy was performed to compare differentially loaded HO shell samples to ensure that samples primarily contained fully formed icosahedral shells rather than aggregates or complex TPI-T1 structures. Cryo-EM imaging showed that all samples contained an abundance of typical shell structures with consistent diameters of 33-35 nm, indicating that the core architecture remained intact even under overloaded conditions (**Figure 3**). The difference in DLS and cryo-EM measurements is expected, as DLS is strongly influenced by flexible surfaces and is biased toward larger species.^40,41^ Visual inspection of the shell interior as captured by cryo-EM revealed more internal density as the supplied TPI concentration increased, demonstrating control over cargo loading (**Figure 3i-o**). In the HT1 TPI 1:3 sample, many particles exhibited additional density extending beyond the shell boundary (**Figures 3h, 3o**). When this extraneous material was included in the measurement, the apparent diameter increased from 35.0 ± 0.13 nm to 42.6 ± 1.04 nm (p = 4.24 × 10^−5^, Student’s t-test). The HT1 TPI 1:3 sample also displayed a noisier background, which may suggest the presence of free, unordered trimer-TPI conjugates. Incomplete encapsulation of the supplied TPI makes it difficult to distinguish between cargo protruding through uncapped vertices and free trimer-TPI conjugants positioned above or below intact shells. In either case, the expanded density mirrors the general trend detected by DLS, suggesting that additional, variably arranged TPI enlarges the particle’s apparent diameter.

**Figure 3.**
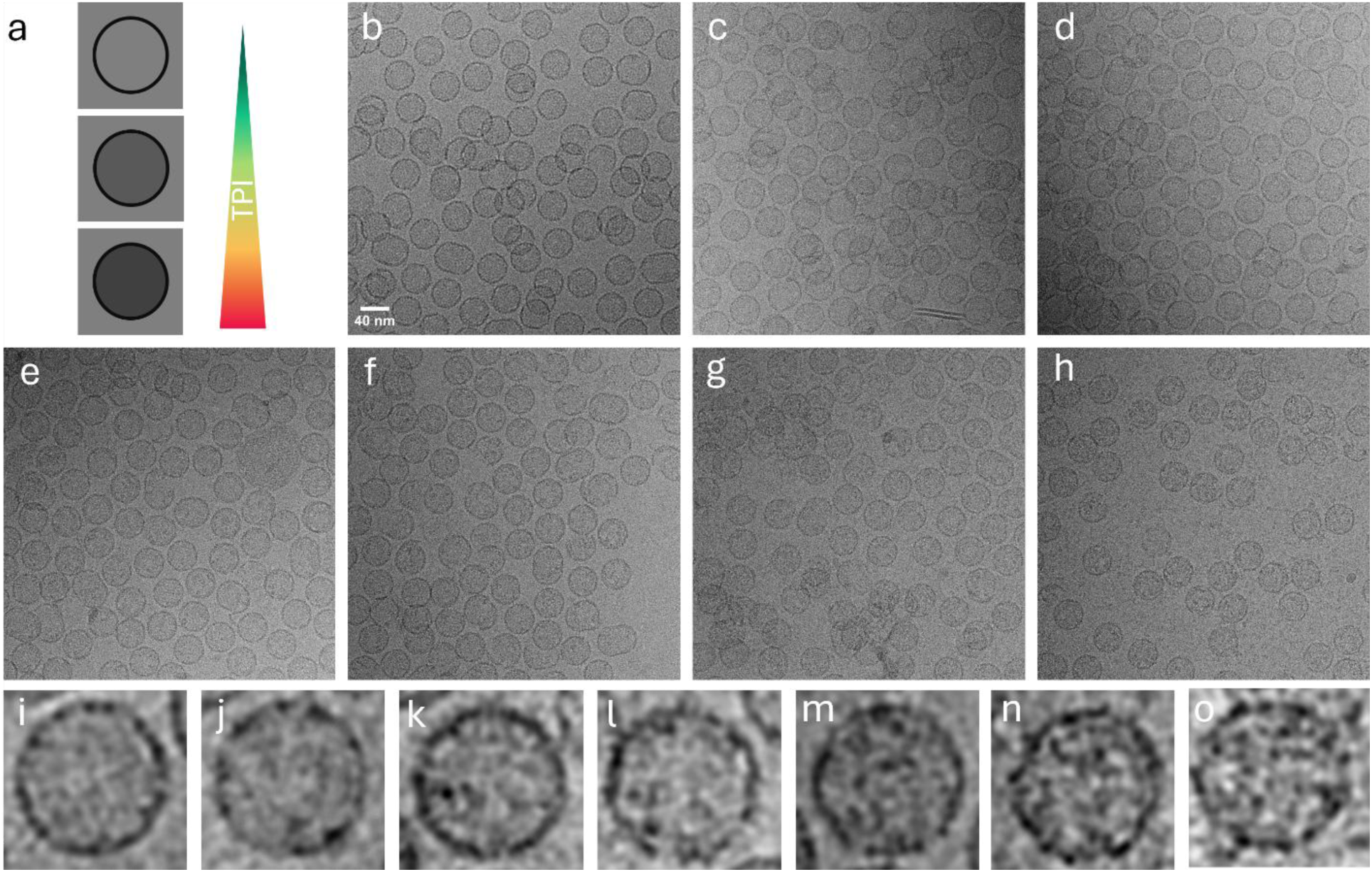
Cyro EM images of variably loaded TPI HO-shells. Cargo remains intact with increasing cargo loads. a) Schematic representation of increasing cargo load. Raw micrographs of BMC shells with **B.** HT1 **C.** HT1 TPI 6:1 **D.** HT1 TPI 3:1 **E.** HT1 TPI 3:2 **F.** HT1 TPI 1:1 **G.** HT1 TPI 1:2 **H.** HT1 TPI 1:3. Single BMC shells with **I.** HT1 **J.** HT1 TPI 6:1 **K.** HT1 TPI 3:1 **L.** HT1 TPI 3:2 **M.** HT1 TPI 1:1 **N.** HT1 TPI 1:2 **O.** HT1 TPI 1:3

### Thermal stability of encapsulated TPI

We next tested thermal stability of encapsulated TPI compared to free TPI by heating the samples and evaluating TPI activity (**Figure 4**). Variably loaded shells and free TPI-SpyCatcher were initially subjected to heating in 10°C increments from 37°C (TPI assay temperature) to 87°C for 1 hour before TPI activity was measured (**Figure S2**). We observed a loss of TPI activity at temperatures above 62°C. For free TPI, loss of function began at 52°C with total loss by 57°C. For encapsulated TPI, normal function was retained up to 52°C and partial function at 62°C. Only at 67°C and above was total loss of activity observed when encapsulated in HT1 shells. The thermal stability assay was repeated with 5°C increments between 37°C and 62°C on TPI and shell samples with T1:TPI ratios of 6:1, 1:1, and 1:3. This experiment showed partial function of free TPI up 52°C (**Figure 4D**). For T1:TPI shells at 6:1 and 1:1 ratios, some activity was retained at 62°C (**Figure 4A and B**). For 1:3 T1:TPI shells, very little activity was observed at 62°C, partial activity at 57°C and 52°C (**Figure 4C)**.

**Figure 4.**
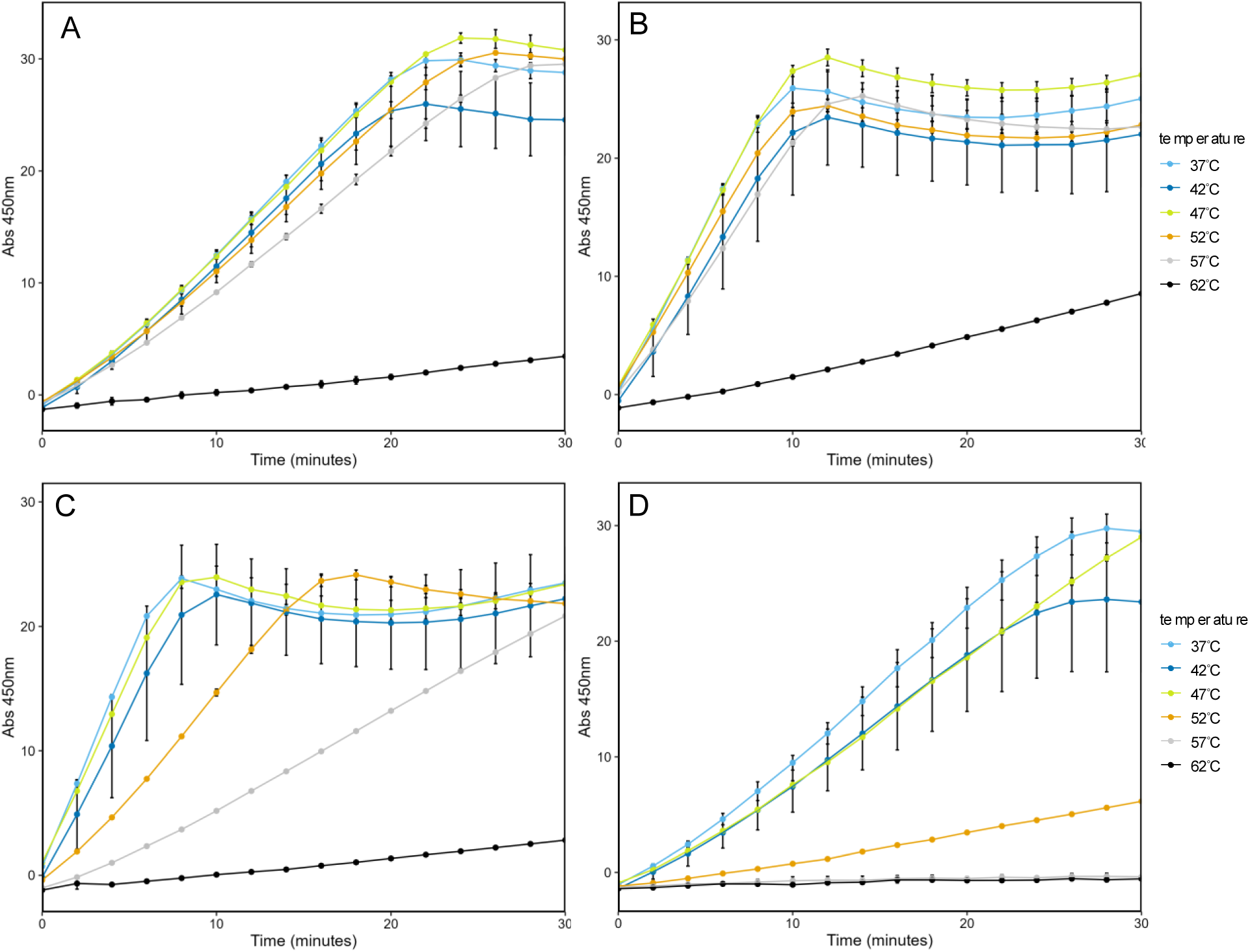
TPI activity of heat-treated HO shells. **A.** HT1 TPI 6:1 **B.** HT1 TPI 1:1 **C.** HT1 TPI 1:3 **D.** TPI. Light blue is 37°C, dark blue is 42°C, yellow is 47°C, orange is 52°C, grey is 57°C, black is 62°C. HT1 samples were done using 238 ng of protein, TPI alone was 2.38 ng.

HT1 TPI 1:3 shells showed a more dramatic loss of activity compared to 6:1 or 1:1 shell samples. This may be due to the presence of a more complex secondary structure of TPI-T1 that could extend outside the shell through the gap caused by the lack of pentamer in these uncapped shells. TPI extending outside the shell may be less protected, leading to a greater loss of activity compared to HT1 TPI 1. DLS results **(Table S1)** may support this and account for the larger measured diameter in these shells.

To ensure that conjugation to the trimer alone was not sufficient for thermal protection we purified T1-TPI and subjected it to the same temperature ranges (**Figure 5**). We observed results intermediate to TPI-SpyCatcher alone and full encapsulation, with some activity retained at 57°C. This may be due to secondary structures formed by the trimer, which can aggregate in solution in the absence of BMC-H, offering some protection to the protein. Regardless, the conjugate is not as stable as fully encapsulated TPI in the HT1 shells, which retains activity up to 62°C.

**Figure 5.**
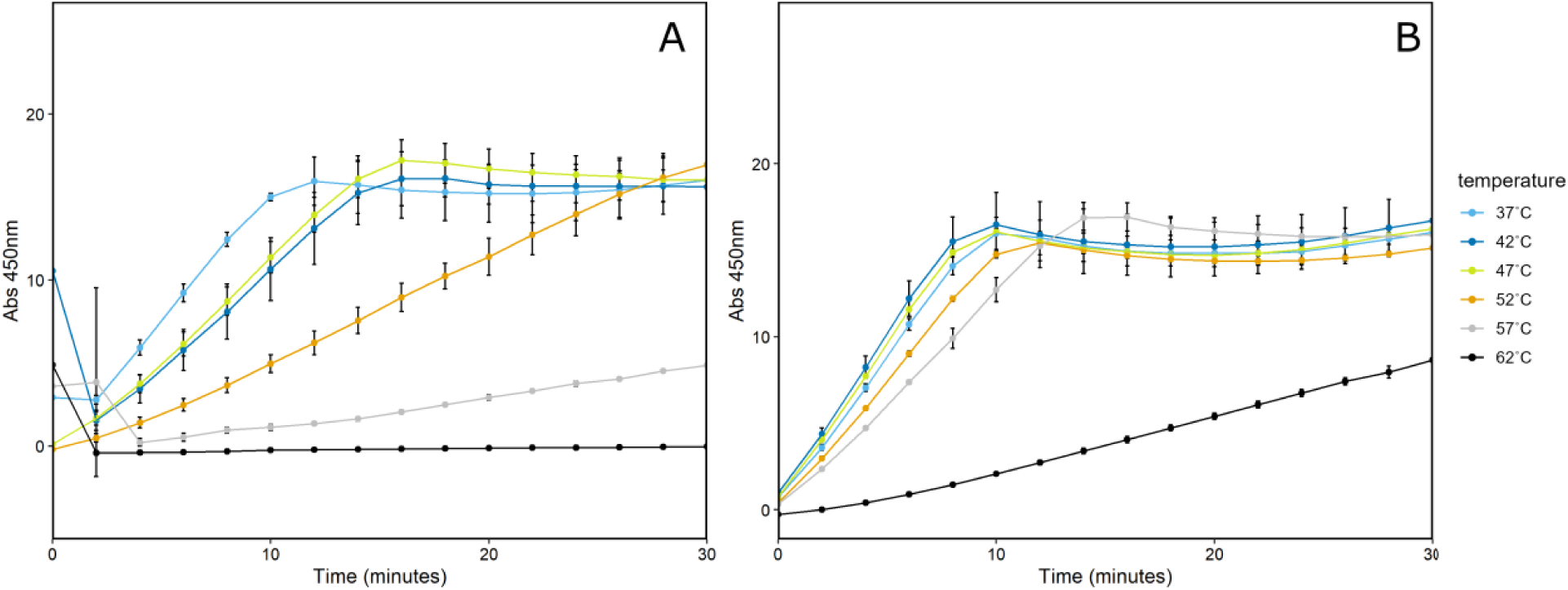
TPI activity of heat treated T1-TPI conjugate. **A.** T1-TPI conjugate purified from SEC treated at a range of temperatures before assay. 238ng of protein. **B.** HT1-TPI 1:1 shells treated at a range of temperatures before assay. 23.8ng of protein. Light blue is 37°C, dark blue is 42°C, yellow is 47°C, orange is 52°C, grey is 57°C, black is 62°C.

### TPI dynamics

The key to understanding the improved TPI stability at elevated temperature within the BMCs are the nanoscale interactions that help confined proteins stay intact under adverse conditions. To obtain molecular-level insight into the thermostability of TPI enzymes inside the BMC shell, molecular dynamics (MD) simulations were employed. MD simulations bridge experimental observations with atomistic descriptions by capturing protein motions and interactions over time. This approach enables the investigation of how confinement and crowding within the BMC shell influence TPI structural stability and dynamics, providing a mechanistic understanding of its enhanced thermal robustness.

Fluctuations at the molecular scale are measured by the root mean squared fluctuation (RMSF), which measures molecular fluctuations around the mean structure. RMSF increases with temperature both when proteins are confined and in solution (**Figures S3 and S4**), since thermal fluctuations increase with temperature. Regions with high RMSF values correspond to flexible segments, typically loops or terminal residues, whereas low RMSF values indicate rigid or structurally constrained regions of the protein. However, the fluctuations increase faster when TPI is in solution alone compared to when it is confined within a BMC. This results in positive peaks when comparing RMSFs between confined and free floating proteins (**Figure 6**). These peaks correspond to protein regions that are more flexible in free solution than when confined in the BMC shell. Region III is particularly interesting, as it is adjacent to glutamate 167 that actively shuttles protons during catalysis,^42,43^ and contains an adjacent isoleucine that displaces water molecules during catalysis.^42^ As temperature increases, the greater dynamics for E167 would likely increase catalysis, up until the point where I172 no longer closes over the substrate and may lead to low affinity and thus poor reactivity. The other three regions identified do not immediately seem related to catalysis but likely indicate protein regions that may unfold first as the protein approaches its melting temperature.

**Figure 6.**
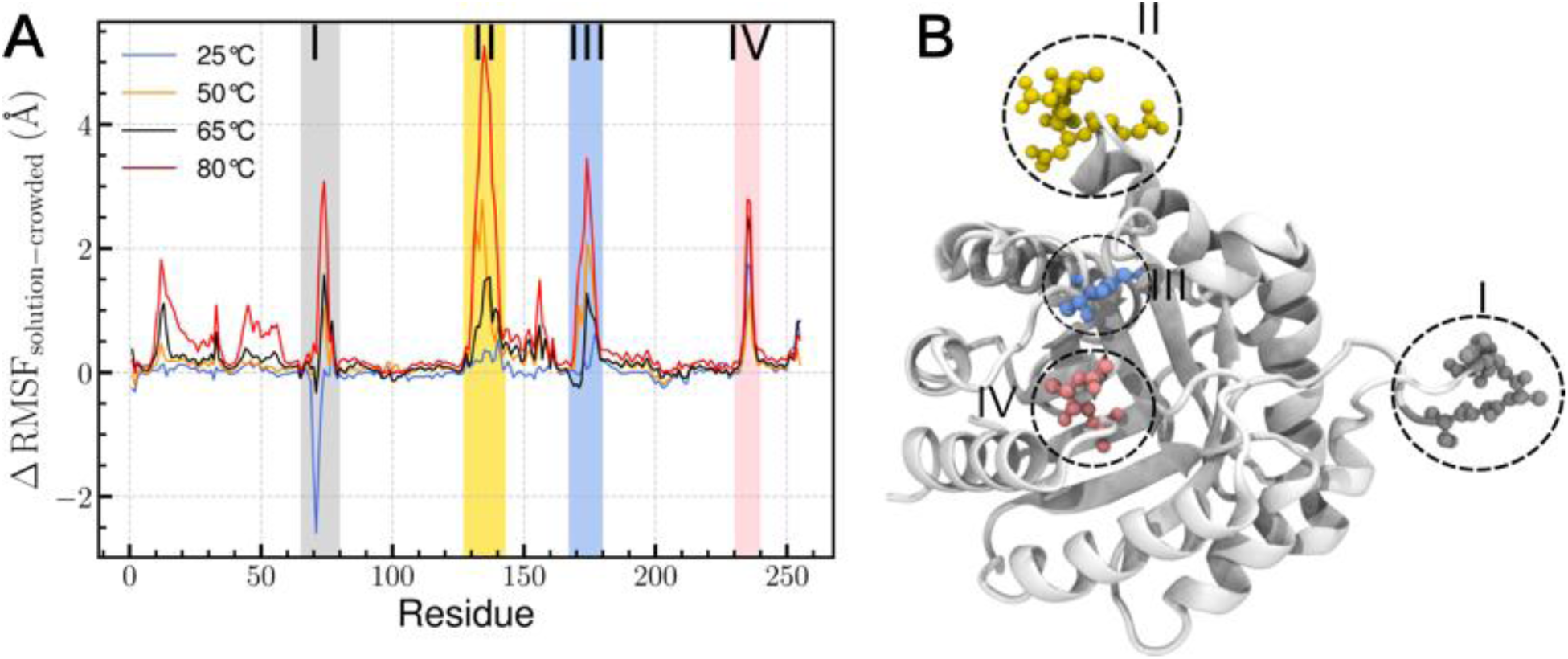
Comparison of TPI dynamics in solution and crowded environment. (A) The difference in RMSF (ΔRMSF) between TPI in solution and in the crowded shell environment is plotted for all residues at multiple temperatures. Regions with ΔRMSF ≥ 2.0 Å are highlighted with colored shading to emphasize regions strongly affected by crowding. Traditional RMSF plots for both TPI in solution and TPI within the shell are provided in Figure S3 and S4. (B) The four highlighted regions are mapped onto the 3D structure of TPI, showing the spatial distribution of residues whose dynamics are most influenced by the crowded environment.

The RMSF are largely identical across most residues when comparing temperature responses and may be underestimates for real conditions due to limited simulation times not allowing the protein to unfold above its melting temperature. We do occasionally see instances where the RMSF is lower in solution than it is in the BMC, particularly for our lowest temperature sample. We have similar levels of sampling between confined simulations (8 dimers for 100 ns) compared to in solution (1 dimer 3 times for 300 ns), but the different simulation lengths may allow for better rotamer sampling in the longer simulation, which may somewhat lower the RMSF for specific residues. However, the broad trend is consistent with other catalysis studies in the literature, where confinement in metal-organic frameworks^44^ or a polymer network^45^ and many other conditions^46^ leads to a lower effective *K*_m_ at elevated temperatures.

While RMSF can zoom into local changes quite clearly, native contacts provide a more global measure of protein structural integrity, reflecting the maintenance of the native fold and its thermal stability^47–49^. In our simulations, TPI in aqueous solution shows a clear decrease in native contacts with increasing temperature (**Figure 7A**), whereas TPI encapsulated within the BMC shell retains native contacts across all tested temperatures (**Figure 7B**). The smaller spread in the crowded system further indicates that confinement and macromolecular crowding suppress thermally induced unfolding motions. As temperature increases, thermal fluctuations progressively disrupt native contacts in solution, whereas in the crowded BMC environment these contacts are retained to a greater extent. Thus, the crowding and confinement delivered by the shell stabilizes the global protein structure under elevated temperature, allowing for accelerated reaction catalysis at higher than usual temperatures.

**Figure 7.**
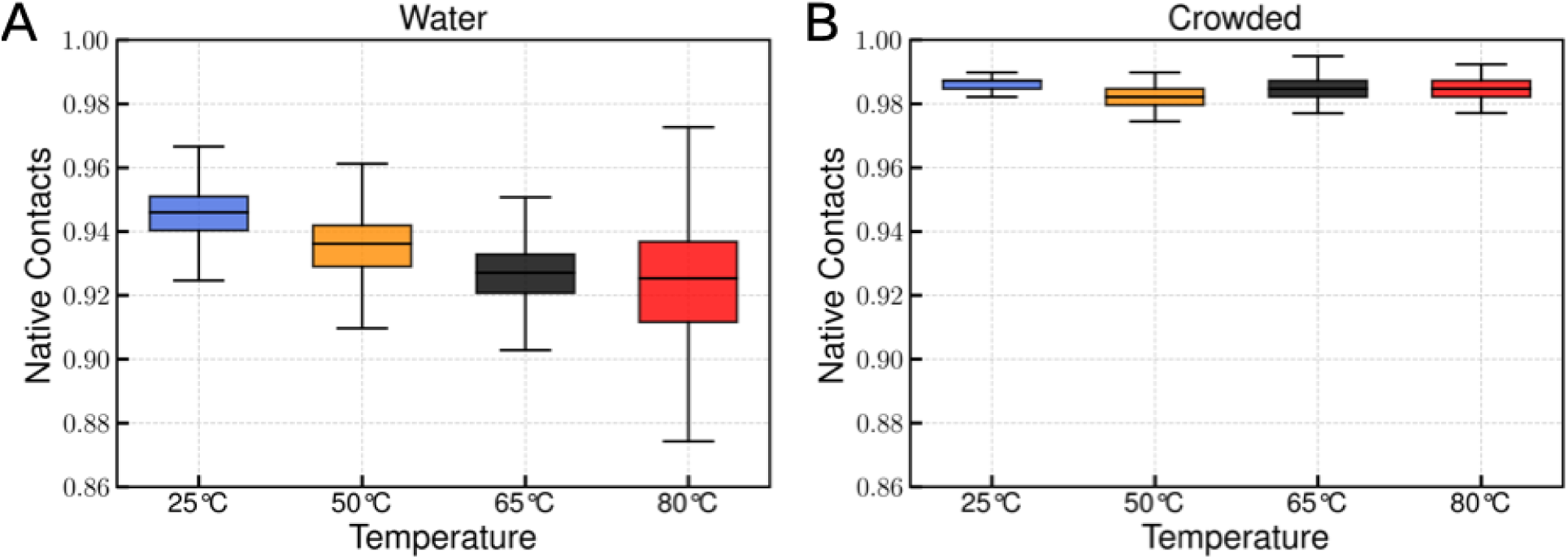
Comparison of TPI structural stability in solution and crowded environments. (A) Fraction of native contacts (Q) for TPI in aqueous solution. (B) Fraction of native contacts (Q) for TPI confined within the crowded BMC shell. Changes in native contact retention reflect the influence of macromolecular crowding on TPI structural stability across temperatures.

## Discussion

By leveraging the IVA toolkit developed by Range et al.,^26^ we reliably produced BMC shells with varying levels of cargo encapsulation. Further, production of TPI-loaded HO shells by IVA was more efficient than previous *in vivo* assembly. Previously, 4 l of *E. coli* culture was needed to yield shells at volumes comparable to a 10 ml IVA. A single preparation of 2 l of culture yielded hexamer sheets sufficient for at least 50 IVA reactions of 10 ml each. Similarly, 2 l of culture produced BMC T1 protein sufficient for at least 15, 10 ml IVA reactions.

Assembly was reliable and simple, but cargo localization initially was not. The SpyTag/Catcher001 system was much slower than the assembly, resulting in underloaded TPI shells or no cargo encapsulation. This necessitated an upgrade to the faster-conjugating SpyCatcher003 sequence on the TPI cargo and SpyTag003 on the trimer. This better matched the speed of conjugation to the speed of shell assembly.^26^ This also allowed us to avoid T1 and T1-TPI aggregation that formed when attempting to pre-conjugate cargo prior to IVA (data not shown).

Here, we demonstrated reliable IVA of HT1 TPI shells with higher activity than comparable samples previously produced by *in vivo* assembly. The increased activity is more impressive when taken together with the lack of background TPI activity in shells without cargo. The “empty” *in vivo* HT1 shells showed 4 times higher TPI activity than empty IVA HT1. We predict IVA will reduce adventitious encapsulation in most situations, assuming effective purification of the target cargo. It is also feasible that switching to SpyCatcher003 instead of SpyCatcher001 could promote increased *in vivo* cargo loading and decrease the incidence of contamination of native proteins. However, given the ease of IVA and lack of background activity in IVA samples, this is likely not worth investigating further.

IVA also improved control of cargo encapsulation for variable loading of HO BMCs. Here, we loaded shells with targeted amounts of TPI and observed TPI activity corresponded well to the expected enzyme loading, with loading down to a T1:TPI ratio of 6:1. Previously, we proposed we had reached the maximal loading possible under *in vivo* assembly of a T1:TPI ratio of 1:1. Here, we increased activity by overloading shells above 1:1 T1:TPI. The 1:2 and 1:3 T1:TPI samples showed an increased abundance of TPI-T1 conjugate and decreased free T1 signal on SDS-page and a corresponding increase in T1-TPI. However, the 1:2 and 1:3 conditions do not show a marked difference in activity, and there may be inhibitory effects of overloading or overcrowding the shells.

To better understand the impact of encapsulation on TPI, we measured thermal stability of encapsulated and free TPI. We observed that confinement of TPI cargo in shells conferred increased thermal stability, with normal activity up to 57°C and partial activity up to 67°C. TPI alone showed normal activity up to 47°C and very low partial activity at 52°C. A 10°C increase in thermal stability of an enzyme is in line with improvements made by direct modification of the enzyme sequence in other studies.^37,38^ Here, we achieved comparable improvement without changing the enzyme primary sequence (other than linking to the SpyCatcher domain). The ability to use BMC shells as a tool for enhancing thermal stability is an exciting finding with the potential to impact many enzymatic reactions if broadly applicable. A great deal of effort in the field has been applied to modifying enzymes or identifying thermophilic enzymes to reach temperatures more suited to industrial settings.^50–54^ The potential applications include enabling enzymes that are poorly suited to higher temperatures to be encapsulated and used more permissively.

Crowding within the BMC shell stabilizes TPI by preserving native contacts and suppressing thermally driven conformational fluctuations that would otherwise interfere with catalysis. In part, the crowded environment within a BMC slows down diffusive motions generally, but particularly near the shell surface ^25^. Indeed, even within 5 nm of the surface, water and metabolite diffusion is noticeably slower. In the smaller model BMC we use here, that corresponds to the entire internal volume for the BMC. Thus, we would anticipate that as the BMC would increase in size, the stabilization due to confinement would decrease somewhat. However, so long as protein loading remains high, we would anticipate that reactivity in large shells would remain high, as the protein diffusion within the shell would also be limited by protein-protein interactions within the shell^55^.

Taken together, these advancements in IVA shell assembly help position the HO BMCs as a potential platform technology, improving enzyme stability and catalysis at elevated temperature through confinement. Future work should be devoted to testing other enzymes under conditions of thermal and other stresses to evaluate how broadly these protections extend.

## Materials and Methods

### Bacterial Strains, Plasmids, and Growth Conditions

Strains and plasmids used in this study are listed in Table 1. pNT003 was generated via PCR linearization of pNT002 to remove the existing TPI-SpyCatcher001 from the N-terminus of TPI. A synthetic gene fragment of TPI-SpyCatcher003 (Twist Bioscience) was then placed in the same position on the N-terminus of TPI using HiFi Master Mix (NEB) with the standard protocol. *E. coli* BL21(DE3) chemically competent cells (Thermo Scientific) were transformed with 15-20 µg/ml of target plasmid(s) using the manufacturer’s standard protocol. After overnight growth, several colonies from the transformation were grown overnight in culture tubes with 5 ml of lysogeny broth (LB) (Miller, Fisher) shaking at 250 rpm at 37°C. Antibiotics were used at the following concentrations: 100 µg/ml ampicillin and/or 50 µg/ml kanamycin.

### Protein purification

2 l of LB were inoculated with 5 ml of preculture at OD_600_=1.0 and induced with 100 mM IPTG or 100 ng/mL anhydrotetracycline (Millipore Sigma). Cells were harvested after 16-18 h of growth by centrifugation at 8000 x g for 10 min at 4°C (Sorvall LYNX 6000, Thermo Fisher) and resuspended in 100 ml of 50 mM Tris pH 8.0, 50 mM NaCl, and 20 mM imidazole by vortexing. 200 µl of DNAse I (Millipore Sigma) and 0.5 tablet of SigmaFast protease inhibitor (Millipore Sigma) were added. Resuspended cells were lysed by passage through a pre-cooled French press at 1,100 PSI twice. Lysate was clarified by centrifugation at 45,000 x g for 30 min at 4°C (Sorvall LYNX 6000, Thermo Fisher). Supernatant was removed and filtered through a 0.22 µm syringe filter. Proteins were purified using a ÄKTA pure fast protein liquid chromatography system (Cytiva) with a 5 ml HisTrap column (Cytiva) and eluted with 50 mM Tris pH 8.0, 50 mM NaCl and 300 mM Imidazole. BMC-T1 or TPI were filtered using a 0.22 µm syringe filter before being further purified using a HiLoad Superdex 200pg SEC column (Cytiva) with buffer containing 50 mM Tris pH 8.0 and 200 mM NaCl to separate proteins from contaminants.

IVA produced BMCs were separated using either a HiLoad Superdex 200 pg (Cytiva) SEC column for 5 ml or Superdex 200 Increase 10/300 GL (Cytiva) for 500 µl assemblies.

### BMC-H Inclusion Body purification

Purification of BMC-H was performed as previously described^26,28^ with the following modifications. BMC-H sheets were resuspended in resuspension buffer containing 8 M urea before storage at -20°C.

### SDS-PAGE

Protein concentrations were measured via bicinchoninic acid (BCA) assay (Thermo Fisher) for assembled BMCs, BMC-T1, and TPI. For BMC-H the presence of urea necessitated the use of absorbance at 280 nm to estimate protein concentration of solubilized sheets. Samples were normalized by dilution in TBS 50/200 buffer to ensure consistent protein loading. 5 μl of each normalized sample was heated at 95°C for 10 min in 5 μl of a mixture of 1 ml 2x loading buffer (Biorad) and 10 μl of 1 M DTT. A mini-PROTEAN tetra cell electrophoresis chamber (Biorad, 1658005EDU) was loaded with 1x TGS buffer. 10 µl of sample was loaded onto a mini-protean TGX stain free gel (Bio-Rad, 4568095) alongside 5 μl of Precision Plus Unstained ladder (Biorad, 1610363). Samples were run at 300 V for 20 min until the dye front moved off the gel. Gels were stained using Coomassie blue for 30 min, then destained in 10% methanol, 10% acetic acid overnight before imaging.

### *In vitro* Assembly

IVA was performed as previously described with the following modifications: Glycerol was omitted from the reaction volume. For a 500 µl IVA 63 µl of 11.87 mg/ml BMC-H, 82 µl of 3.27 mg/ml BMC-T1, and between 0-270 ul of 1.47 µg/ml TPI-SpyCatcher003 was used. For large scale IVA these volumes were increased 10-fold.

### Triose phosphate isomerase activity assay

Tpi activity was measured using Triose Phosphate Isomerase (TPI) Activity Assay Kit (Colorimetric) (Abcam, ab197001) as described in the product manual. Protein samples were standardized to total protein using the BCA assay described above. For temperature gradient experiments, samples were prepared in Assay Buffer II from the above kit to a final volume of 50 ul in PCR tubes. Samples were then heated for one hour before cooling for 15 minutes then transferred to a 96-well plate for the TPI assay.

### Dynamic light scattering analysis

Dynamic light scattering was performed on a Wyatt DynaPro (Nanostar). 10 μL of the HO shell samples were centrifuged for 5 min at 13,000 x g before being loaded into 1×1×10 mm cuvette. Samples were scanned 20 times with 5 second acquisitions. This was repeated three times on each sample to measure shell diameter.

### Cryo-electron microscopy

A 4 µl aliquot of assembled shells was loaded onto a R 2/2 Cu 200 Quantfoil grid that had been freshly glow-discharged for 45 seconds using a Pelco Easiglow. After a wait time of 10 seconds at 4°C and 100% humidity, the grid was blotted onto filter paper using a blot force of 1 for 4 seconds then plunge-frozen into liquid ethane using a Vitrobot Mark IV. Data were collected at the Michigan State University RTSF Cryo-EM facility on a Talos Arctica operating at 200 keV and equipped with a Falcon 4i camera. Micrographs were collected at 130,000x nominal magnification (0.886 Å/pixel) by recording 1 frame over 6 seconds for a total dose of 22.8 e/Å^2^.

### Computational Methods: Aqueous TPI System

To evaluate the mechanism for improved thermostability under confinement, TPI was simulated in both aqueous solution (Fig. 1A, B) and within a BMC shell (Fig. 1C-E) at four temperatures (25°C , 50°C , 65°C, and 80°C). The simulation protocols were kept consistent across both systems to allow direct comparison of the protein dynamics both in a confined space and in free solution. While obtaining initial coordinates for the protein component was straightforward using existing PDB structures, a key challenge was filling the initially hollow shell. This required not only balancing water across the shell boundary but also efficiently packing multiple TPI dimers inside the BMC shell.

For the aqueous system, a single TPI dimer (PDB ID: 4K6A)^56^ was placed in a 104 Å long cubic water box using the solvate plugin within VMD^57^, ensuring sufficient padding to ensure complete solvation for the dimer structure (**Figure 8A, B**). Production molecular dynamics simulations were carried out for 300 ns from this solvated starting point for three independent replicas, resulting in 900 ns of cumulative sampling per temperature. These trajectories provided a robust baseline for assessing the structural and dynamical properties of TPI in solution, in the absence of crowding.

**Figure 8.**
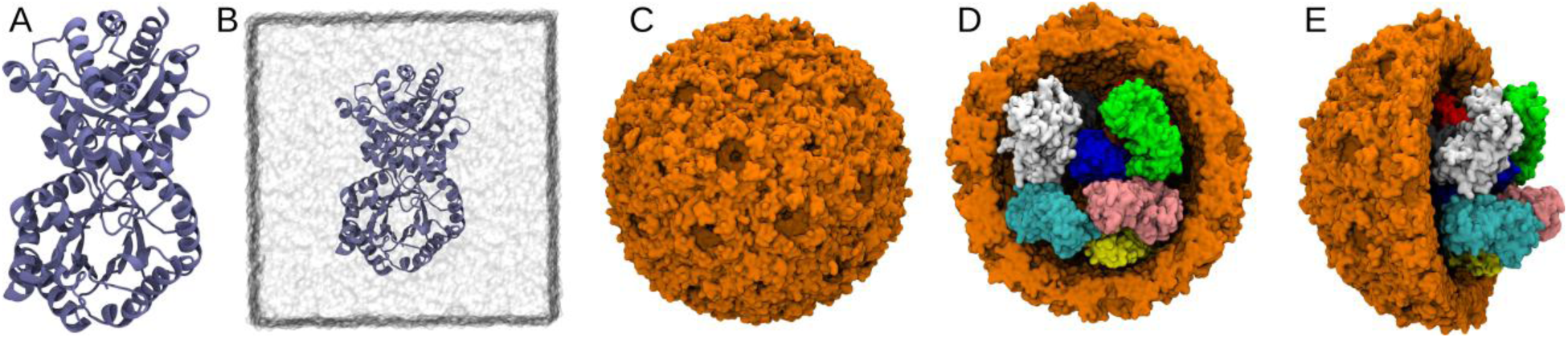
Pictorial representation of TPI in solution and in a crowded BMC environment. Panels A and B show the 3D atomistic representation of TPI, with panel B highlighting TPI solvated in a water box. Panel C illustrates the complete HP shell. Panels D and E show the front and side views of the HP shell encapsulating TPI. Water molecules and ions are omitted for clarity.

### Computational Methods: Shell Preparation and TPI Packing

For modeling the crowded environment, we economized on the shell size compared to the full HO shell that encapsulated the protein in experiment. Instead of the full HTP shell, we encapsulated TPI dimers within an analogous HP shell (PDB ID: 6OWG)^58^ from another organism, which has a smaller diameter and thus a smaller simulation footprint. To determine a physiologically relevant number of TPI molecules for encapsulation, we considered the typical protein volume fraction in bacterial cytoplasm, ϕ ≈ 0.2, which is similar to prior estimates for protein loading in BMC shells ^23,59^. When we assume that the shell interior and TPI are spherical, we needed 9 TPI dimers to occupy 20% of the internal shell volume. The 9 TPI dimers were manually placed within the HP shell to avoid contacts with each other and with the shell surface. This approach ensures that the protein occupancy inside the shell is consistent with physiological crowding constraints (**Figure 8, C-E**). Production simulations were then performed for 100 ns at each temperature.

### Computational Methods: Equilibrium Simulation Protocol

To investigate the atomic-level dynamics of TPI, we performed classical MD simulations using the CHARMM36m force field^60^. Explicit solvation was modeled with the TIP3P water model^61^ and the charges were neutralized by adding Na^+^ and Cl^−^ ions such that the resulting concentration is around 0.15 M. All simulations were carried out in NAMD 3.1 alpha1 using the GPU-resident integrator to maximize performance^62^. Pressure was maintained at 1 atm with the Langevin piston method^63^, and temperature was controlled using a Langevin thermostat with a damping coefficient of 1 ps^−1^. Though this damping slows diffusion relative to water dynamics in other integrators, it provides stable temperature control across all systems. Hydrogen bond lengths were constrained using SETTLE, enabling a 2 fs integration timestep^64^. Long range electrostatics were computed using particle-mesh Ewald (PME) with a grid spacing of 1.2 Å, while short-range nonbonded interactions employed a 12 Å cutoff with a 10 Å switching distance^65,66^. Each system was first energy minimized for 5000 steps using conjugate gradient minimization^67^. After minimization, velocities were reinitialized at 298K and the system was equilibrated for 1 ns prior to production simulations.

### Computational Methods: Trajectory Analysis

Trajectory analysis was performed using a combination of built-in and custom Python scripts in VMD^57^, leveraging the NumPy and SciPy libraries^68,69^, for numerically intensive calculations. In the crowded system, one of the nine TPI dimers exhibited steric clashes not identified during placement with the interior of the BMC shell, leading to partial misfolding. This dimer was therefore excluded from all subsequent structural and dynamical analyses to ensure accurate characterization of TPI dynamics and structure. Analyses included calculation of root-mean-square fluctuations (RMSF), native contacts, and other measures of protein conformational dynamics, providing insight into the effects of crowding on TPI stability and flexibility. Since TPI is a homodimer, we choose to present the results by combining statistics for each of the constituent monomers, increasing our statistical power.

### Computational Methods: Root-Mean-Square Fluctuations (RMSF)

To quantify the local flexibility of TPI residues, root-mean-square fluctuations (RMSF) of Cα atoms were calculated from entire production run. For each protein, the RMSF was computed as-

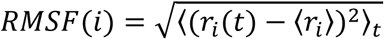

where r_i_(t) is the position of the i^th^ Cα atom at time t, and ⟨ri⟩ is the time-averaged position of that atom over the trajectory. For the aqueous system, RMSF values were averaged over three independent replicas to ensure convergence. In the crowded system, RMSF was computed for the eight TPI dimers that did not exhibit steric clashes. Comparisons between RMSF profiles in solution and in the crowded shell provide a measure of how macromolecular crowding affects the flexibility of individual residues. High RMSF values correspond to flexible regions, often loops or terminal residues, while low RMSF values indicate rigid or structurally constrained regions of the protein.

### Computational Methods: Native Contacts

Native contacts were used to quantify the structural integrity and thermal stability of TPI throughout the simulations, as the preservation of native contacts reflects the extent to which the folded state is maintained under thermal stress. The reference native structure from the PDB was used to define the existing native contacts that we will compare to. Two residues i and j were considered to form a native contact if the distance between their heavy atoms (Cα) was less than 4.5Å in the reference structure and if |i − j| > 3, thereby excluding trivial local contacts.

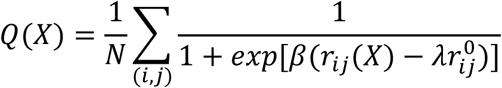

where the sum runs over the N native contact pairs (i, j), r_ij_(*X*) is the distance between 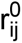 structure. The parameter β = 5Å^−1^controls the sharpness of the contact transition, while the scaling factor λ = 1.8 defines the cutoff for contact formation and accounts for thermal fluctuations.

Native contacts were calculated for all temperatures in both aqueous and crowded systems. Since TPI is a homodimer, native contact analysis was performed on individual monomers, and results are reported for monomers only for clarity.

## Abbreviations

BMCs: bacterial microcompartments
BMC-H: hexamer
BMC-P: pentameter
BMC-T1: trimer
DLS: dynamic light scattering
FPLC: fast protein liquid chromatography
HO: *Haliangium ochraceum*
HT1: minimal shell bacterial microcompartment
HT1T2T3: full shell bacterial microcompartment
IVA: *in vitro* assembly
MD: molecular dynamics
RMSF: root mean squared fluctuation
SEC: size exclusion chromatography
TPI: triose phosphate isomerase

## Acknowledgements

Research was supported as part of the Center for Catalysis in Biomimetic Confinement, an Energy Frontier Research Center funded by the U.S. Department of Energy (DOE), Office of Science, Basic Energy Sciences (BES), under award DE-SC0023395. This work was supported in part through computational resources and services provided by the Institute for Cyber-Enabled Research at Michigan State University. This research used resources of the National Energy Research Scientific Computing Center (NERSC), a Department of Energy User Facility using NERSC award BES-ERCAP m4325.

