## Supplementary figures and table for "Encapsulation in a bacterial microcompartment shell improves thermal stability of a glycolytic enzyme"

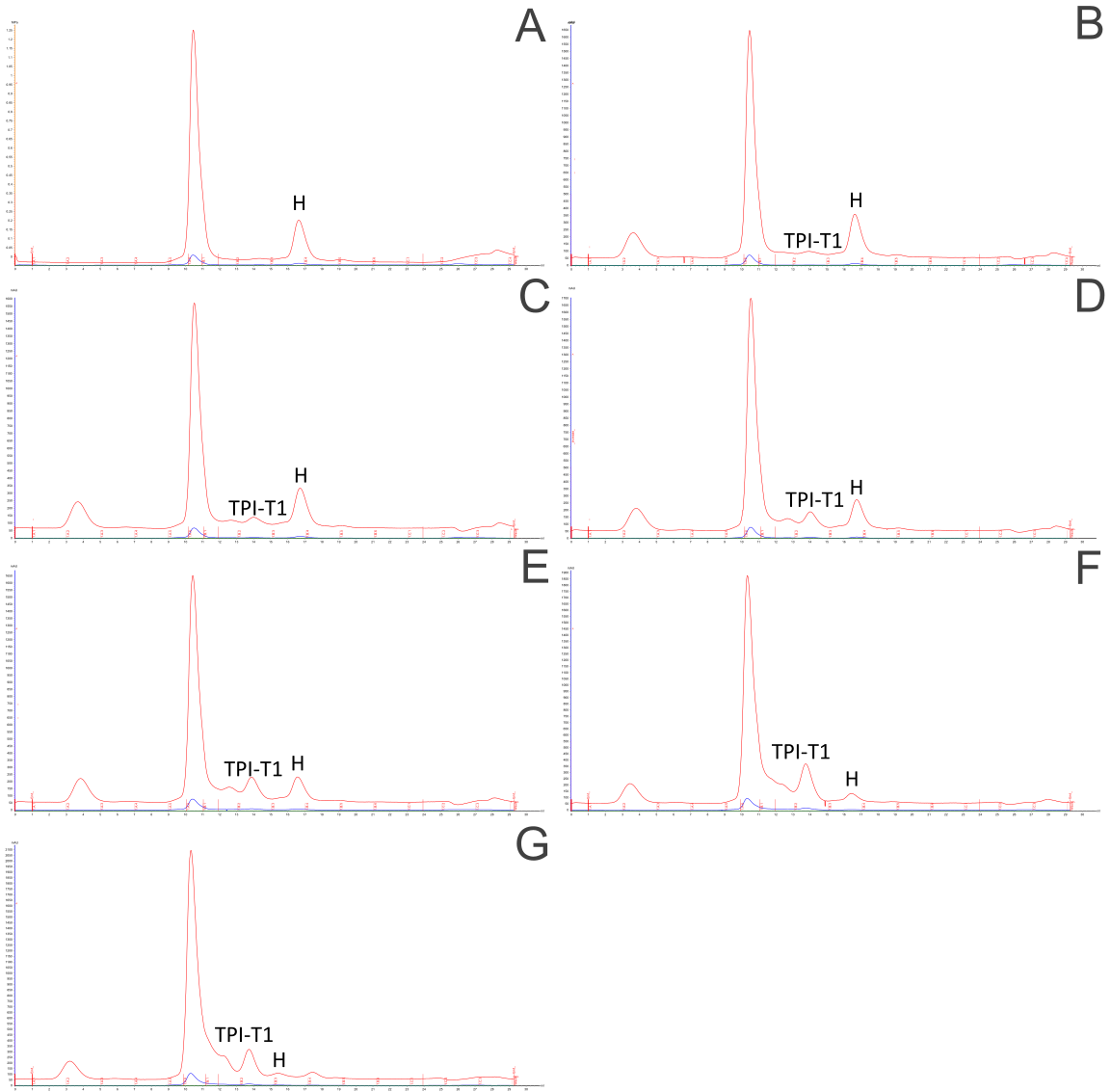

**Figure S1. Size exclusion chromatograms of variably loaded HT1 shells. A.** HT1 **B.** HT1 TPI 6:1 **C.** HT1 TPI 3:1 **D.** HT1 TPI 3:2 **E.** HT1 TPI 1:1 **F.** HT1 TPI 1:2 **G.** HT1 TPI 1:3

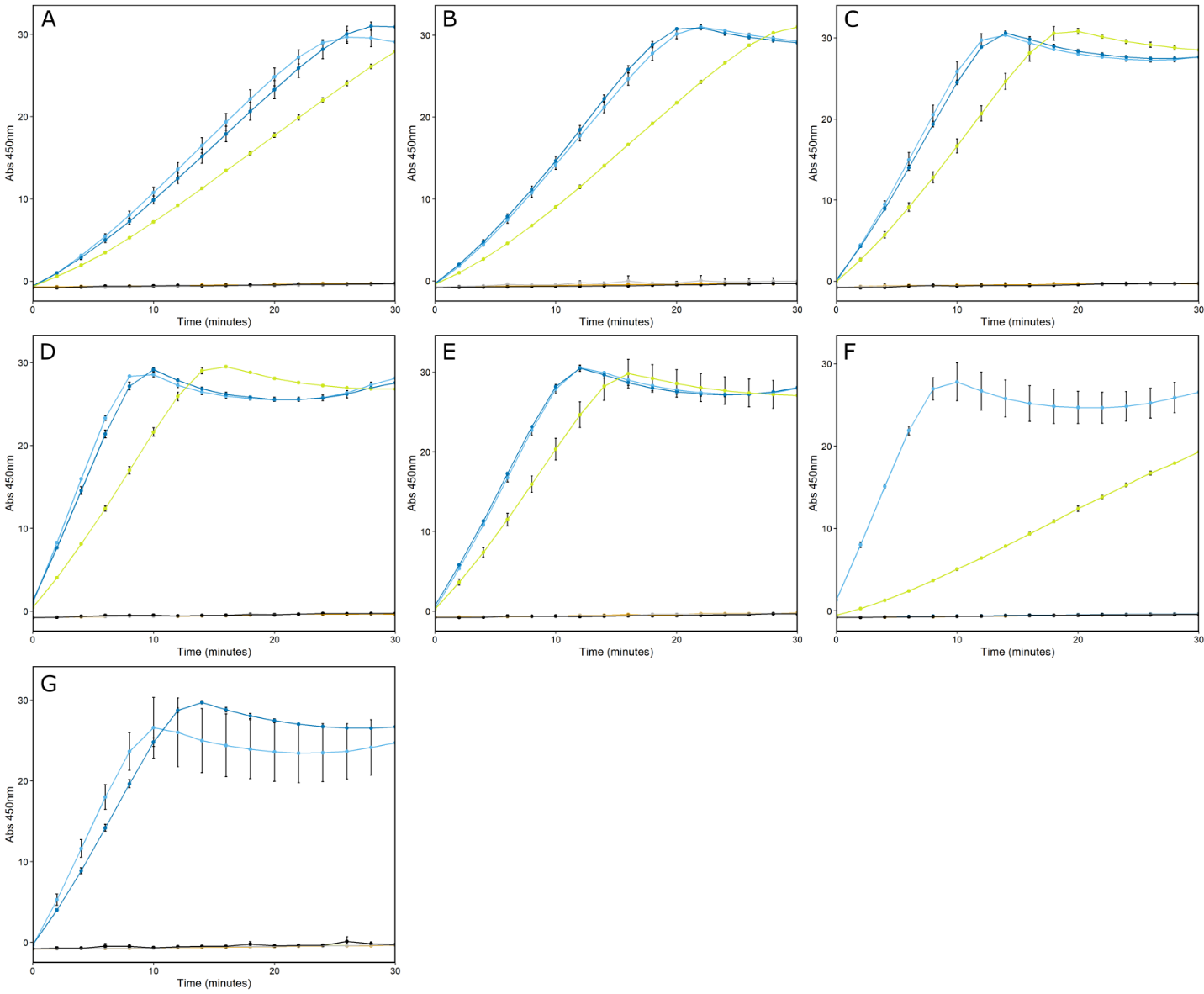
**Figure S2. TPI activity of heat-treated HO shells. A.** HT1 TPI 6:1 **B.** HT1 TPI 3:1 **C.** HT1 TPI 3:2 **D.** HT1 TPI 1:1 **E.** HT1 TPI 1:2 **F.** HT1 TPI 1:3 **G.** TPI. Light blue is 37°C, dark blue is 47°C, yellow is 57°C, orange is 67°C, grey is 77°C, black is 87°C.

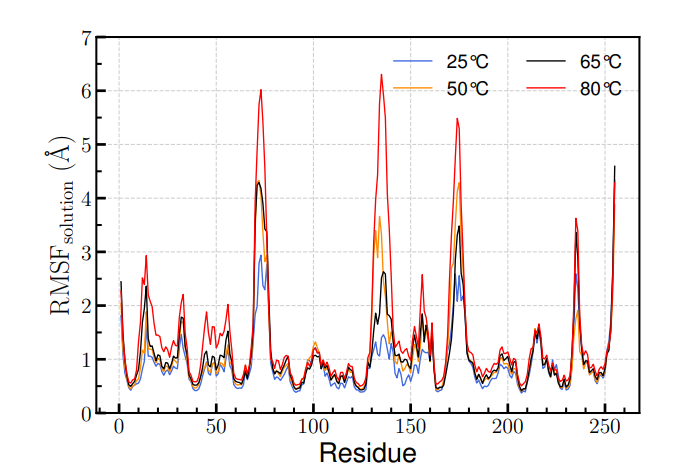

**Figure S3. RMSF of TPI in aqueous system.**

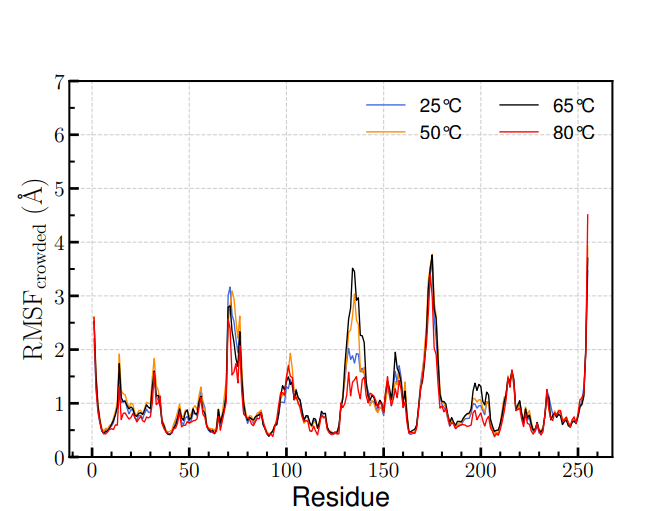

**Figure S4. RMSF of TPI in crowded system.**

**TableS1.** Dynamic Light Scattering analysis of shell samples. Each shell average diameter is an average of 3 reads.

| Shell | average  diameter (nm) | stdev |
| --- | --- | --- |
| HT1 | 44.22 | 0.57 |
| HT1 TPI 6:1 | 46.05 | 0.38 |
| HT1 TPI 3:1 | 45.96 | 0.23 |
| HT1 TPI 3:2 | 46.65 | 0.22 |
| HT1 TPI 1:1 | 47.29 | 0.11 |
| HT1 TPI 1:2 | 48.81 | 0.21 |
| HT1 TPI 1:3 | 57.98 | 1.00 |
